# Computational Hot-Spot Analysis of the SARS-CoV-2 Receptor Binding Domain / ACE2 Complex

**DOI:** 10.1101/2020.08.06.240333

**Authors:** Pedro A. Rosario, Brian R. McNaughton

## Abstract

Infection and replication of SARS CoV-2 (the virus that causes COVID-19) requires entry to the interior of host cells. In humans, a Protein-Protein Interaction (PPI) between the SARS CoV-2 Receptor-Binding Domain (RBD) and the extracellular peptidase domain of ACE2, on the surface of cells in the lower respiratory tract, is an initial step in the entry pathway. Inhibition of the SARS CoV-2 RBD / ACE2 PPI is currently being evaluated as a target for therapeutic and/or prophylactic intervention. However, relatively little is known about the molecular underpinnings of this complex. Employing multiple computational platforms, we predicted ‘hot-spot’ residues in a positive control PPI (PMI / MDM2) and the CoV-2 RBD/ACE2 complex. Computational alanine scanning mutagenesis was performed to predict changes in Gibbs’ free energy that are associated with mutating residues at the positive control (PMI/MDM2) or SARS RBD/ACE2 binding interface to alanine. Additionally, we used the Adaptive Poisson-Boltzmann Solver to calculate macromolecular electrostatic surfaces at the interface of the positive control PPI and SARS CoV-2 / ACE2 PPI. Collectively, this study illuminates predicted hot-spot residues, and clusters, at the SARS CoV-2 RBD / ACE2 binding interface, potentially guiding the development of reagents capable of disrupting this complex and halting COVID-19.

## Introduction

The novel coronavirus SARS-CoV-2 causes Coronavirus disease 19 (COVID-19), an established global health crisis.^1, 2^ The COVID-19 pandemic continues to take a significant toll on the global economy and healthcare infrastructure, and is attributed to more than 700,000 deaths worldwide; over 160,000 in the United States of America alone. The number of infections, and resulting COVID-19-related deaths continues to grow at a staggering rate. Older individuals, and those with pre-existing conditions, are at the highest risk of death as a result of deleterious consequences related to SARS-CoV-2 infection. Moreover, negative outcomes associated with SARS-CoV-2 infection disproportionately correlate with minorities, likely due to a number of factors.^3^ The magnitude of the COVID-19 pandemic has triggered a global response from researchers to identify therapeutics and/or prophylaxis capable of inhibiting, or slowing, infection and propagation of this deadly disease.

SARS-CoV-2 must enter host cells to replicate. Previous studies on SARS-CoV, which is closely related to SARS-CoV-2, provided insight into the mechanism of cell entry. As an early step in the cell entry mechanism, the viral spike protein (S) on the surface of SARS-CoV binds the extracellular protease domain (PD) of angiotensin-converting enzyme 2 (ACE2), a protein on the surface of multiple cells, including cells in the lower respiratory tract.^4–7^ This recognition event is followed by membrane fusion leading to host cell entry. It was recently shown that SARS-CoV-2 also relies on an interaction between its spike protein and ACE2 to gain entry to the interior of human cells. Cryo-electron microscopy (cryo-EM) structural analysis of the complex involving the SARS-CoV-2 spike protein Receptor Binding Domain (RBD) and human ACE2 was recently reported (PDB code: 6M17), with an overall resolution of 2.9 Å and local resolution of 3.5 Å at the RBD / ACE2 binding interface (**Figure 1a**).^8^

**Figure 1.**
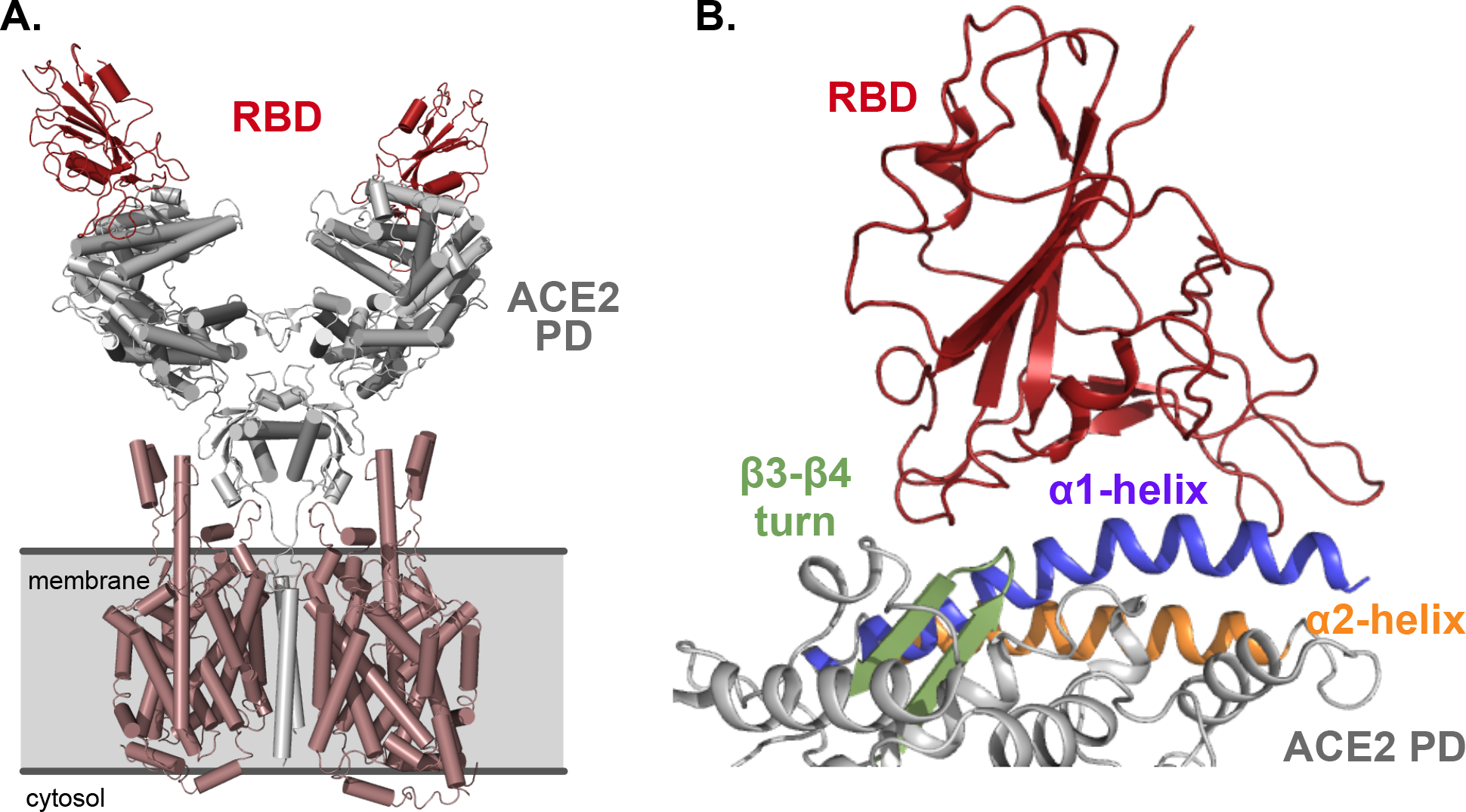
(**A**) Structure of the SARS-CoV-2 RBD / full-length ACE2 complex, colored by subunits. Extacellular ACE2 protease domain (PD) is colored grey; SARS-CoV-2 RBD is colored red. Cell membrane is depicted with a grey rectangle. (**B**) Interface of the SARS-CoV-2 RBD / ACE2 complex. SARS-CoV-2 RBD is colored red; ACE2 α1-helix is colored purple; ACE2 α2-helix is colored orange; ACE2 β3-β4 loop is colored green.

Given the important role the Protein-Protein Interaction (PPI) between SARS-CoV-2 RBD and ACE2 plays in host cell entry, inhibition of this PPI has received attention as a target for therapeutic and/or prophylactic intervention.^9, 10^ Regions on ACE2 closest to SARS-CoV-2 include the α1-helix (**Figure 1b,** blue) α2-helix (**Figure 1b**, orange) and the β3-β4 loop (**Figure 1b**, green). Researchers have shown that a 23 residue peptide, consisting of ACE2 α1-helix sequence (I21-S43) tightly binds SARS-CoV-2 (*K_D_* ~50 nM).^10^ However, relatively little is known about the molecular underpinnings that stabilize the SARS-CoV-2 / ACE2 complex. In an era when academic laboratories have been closed, or severely limited due to the COVID-19 pandemic, we sought to use multiple computational platforms, with an emphasis on servers that are available online to all researchers, to study the SARS-CoV-2 RBD / ACE2 complex examine the electrostatics and topology of the interface, and identify “hot-spots” – amino acids, or clusters thereof, that significantly contribute to the binding free energy of this therapeutically-relevant PPI.

## Results

### KFC-2 Hot-Spot Prediction

We used the <u>K</u>nowledge-based <u>F</u>ADE and <u>C</u>ontacts 2 (KFC-2), a free server that compares favorably to Robetta, FOLDEF, HotPoint, and MINERVA, to predict hot-spot residues and/or clusters at the SARS-CoV-2/ACE2 binding interface.^11, 12^ As a positive control in our study, we performed an analogous examination of the PMI / MDM2 complex (PDB code: 3EQS, **Figure 1a**), which is well studied.^13^ PDB files for these two complexes were loaded into the server, which revealed hot-spot residues at each interface. KFC accurately predicted hot-spot residues on the PMI peptide, which have been experimentally validated (**Figure 2a**).^13^ Specifically, residues F3, Y6, W7, and L10 were identified as hot-spot residues on the PMI peptide. Complimentary hot-spot residues L54, V75, V93, and I99 were identified as hot-spot residues on MDM2 (**Figure 2a-b**; hot-spot scores are shown in **Figure S1, Supporting Information**). Identical analysis of the SARS-CoV-2 RBD / ACE2 complex illuminated five residues on both ACE2 and SARS-CoV-2 RBD that were characterized as hot-spot residues (**Figure 2c-d**; values shown in **Figure S2, Supporting Information**). Specifically, L455, Y489, Q498, T500 and Y505 were identified on SARS-CoV-2 RBD and T27, H34, D38, Y41, and K353 were identified on ACE2. For both complexes, KFC-a values were used, as these have been shown to have superior predictability outcomes compared to KFC-b values.^12^

**Figure 2.**
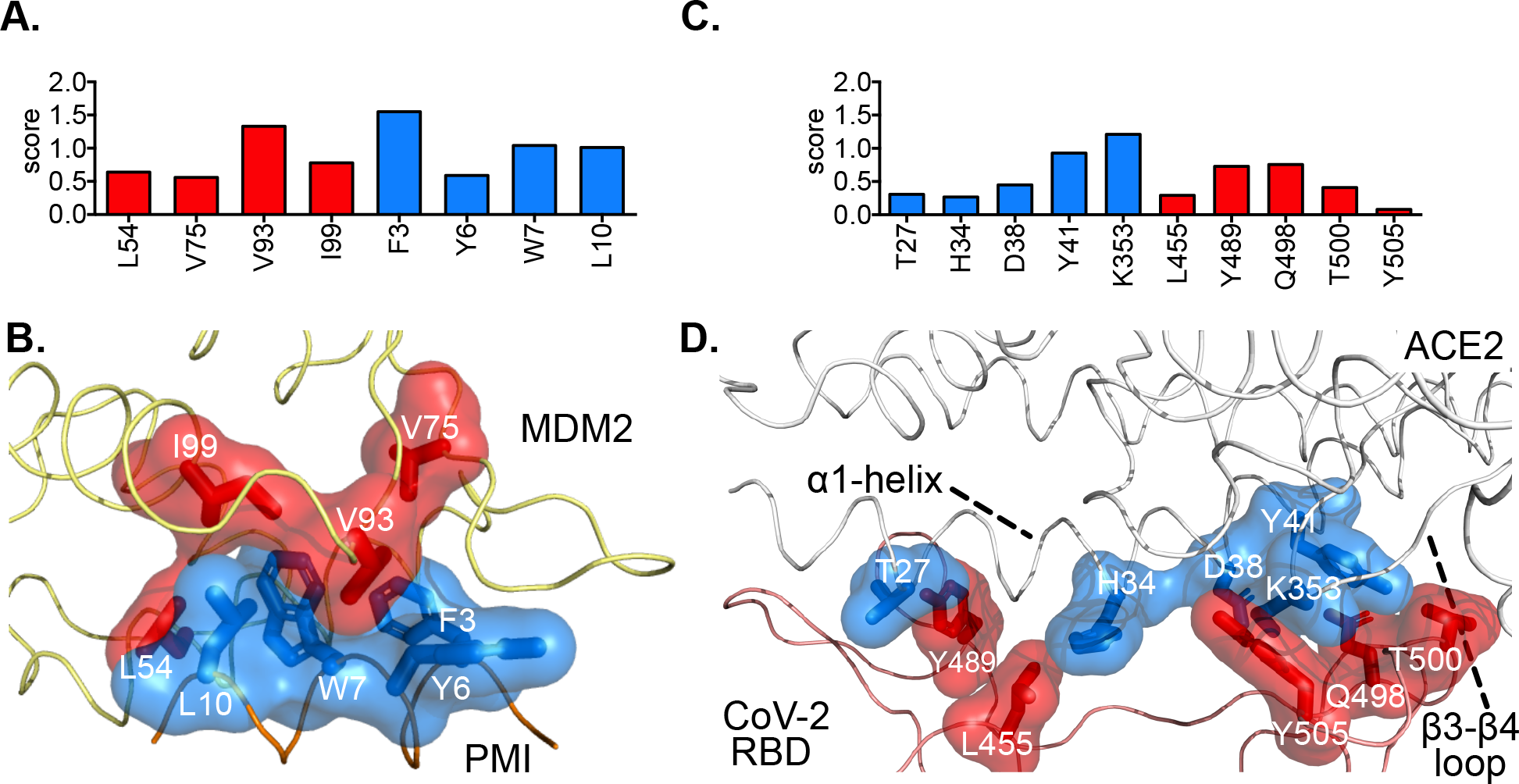
(**A**) Hot-spot residue scores predicted by KFC-2 for the PPI involving PMI and MDM2. (**B**) Structure of the PMI / MDM2 interface, with predicted PMI or MDM2 hot-spot residues highlighted in blue or red, respectively. (**C**) Hot-spot residue scores predicted by KFC-2 for the PPI involving SARS-CoV-2 RBD and ACE2. (**D**) Structure of the SARS-CoV-2 RBD / ACE2 interface, with predicted ACE2 or SARS-CoV-2 RBD hot-spot residues highlighted in blue or red, respectively.

Satisfyingly, four of the five predicted hot-spot residues on ACE2 (T27, H34, D38, and Y41) are found on the α1-helix (**Figure 1b**, blue), and researchers have shown that this helix alone (as a chemically synthesized 23 residue peptide) binds SARS-CoV-2 RBD with a dissociation constant of ~50 nM.^10^ Lysine 353 was also identified as a hot-spot residue, and is on the β3-β4 loop that neighbors the α1-helix (**Figure 2d**). When paired with experimental results, demonstrating high affinity between ACE2 α1-helix and SARS-CoV-2 RBD, these results suggest that KFC-2 correctly identified residues at the interface critical to stability of the complex.

### BAlaS Computational Alanine Scanning Mutagenesis

We next used BAlaS^14, 15^, a computational alanine scanning mutagenesis server, to predict the change in free energy (ΔΔG) associated with mutating interface residues to alanine. Our analysis focused on residues within 13 Å from the binding interface (set as default) and residues plotted in **Figure 3** are limited to those with predicted ΔΔG > 4 kJ/mol. BAlaS correctly identified hot-spot residues F3 (4.6 kcal/mol), Y6 (2.4 kcal/mol), W7 (5.2 kcal/mol), and L10 (2.1 kcal/mol) on PMI. Residues L54 (4.6 kcal/mol), Y67 (2.4 kcal/mol), Q72 (1.7 kcal/mol), H96(1.1 kcal/mol) and V93 (1.5 kcal/mol) were also identified as primary contributors to the binding free energy of the complex (**Figure 3a-b**).

**Figure 3.**
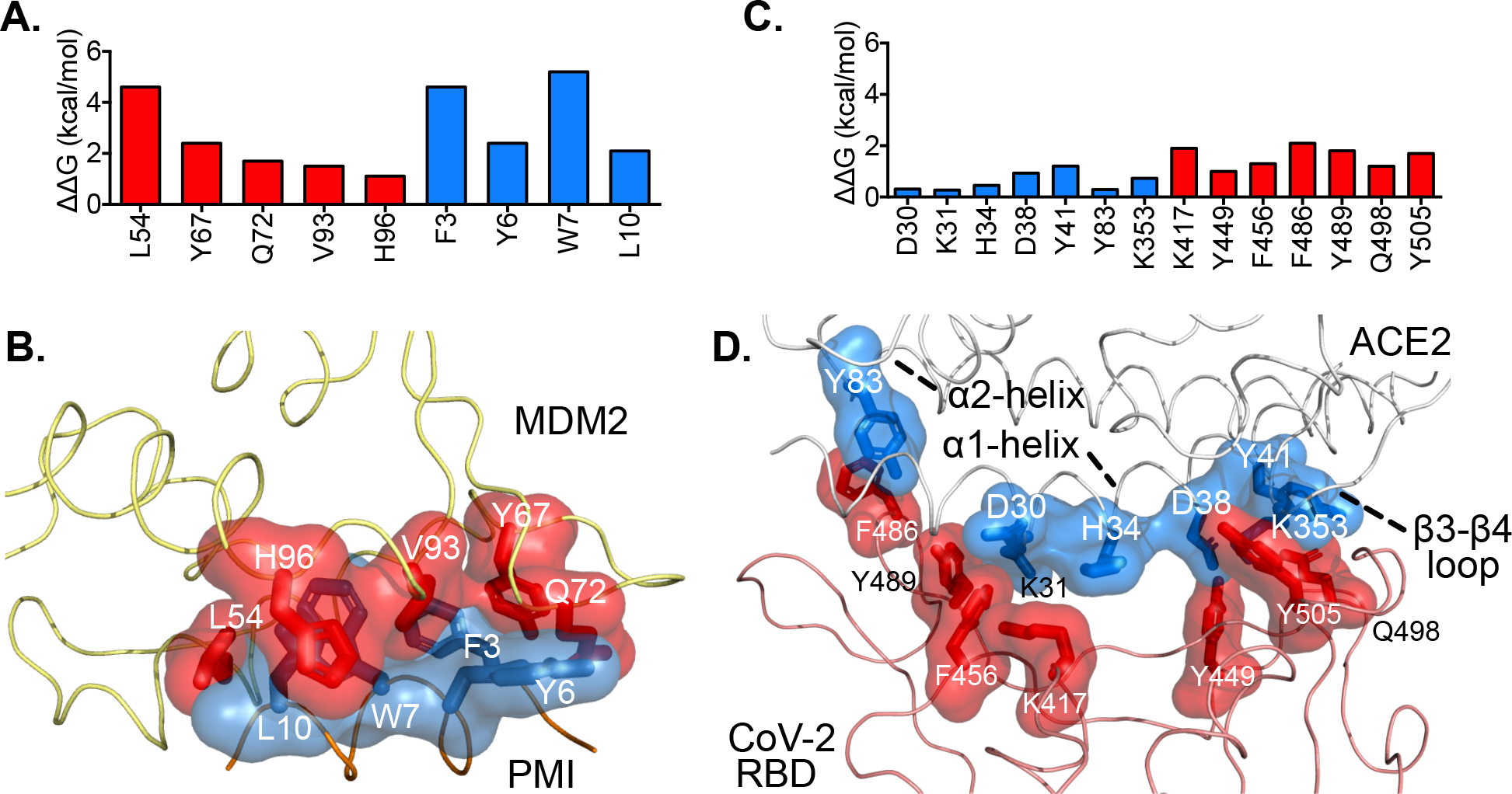
**(A)** Computational alanine scanning mutagenesis data for the PMI / MDM2 complex. (**B**) Residues in the PMI / MDM2 complex that are predicted to contribute significantly to the free energy of the complex. (**C**) Computational alanine scanning mutagenesis data for the SARS-CoV-2 RBD / ACE2 complex. (**D**) Residues in the SARS-CoV-2 RBD / ACE2 complex that are predicted to contribute significantly to the free energy of the complex.

BAlaS analysis of the SARS-CoV-2/ACE2 complex resulted in the prediction of seven residues on ACE2 that contribute to the binding free energy of the complex (**Figure 3c-d**). The following residues, and associated changes in free energy, were identified: D30 (0.3 kcal/mol), K31 (0.3 kcal/mol), H34 (0.5 kcal/mol), D38 (0.9 kcal/mol), Y41 (1.2 kcal/mol), Y83 (0.3 kcal/mol), and K353 (0.7 kcal/mol). ACE2 residues, and associated predicted changes in binding free energy as a result of alanine mutagenesis are as follows: K417 (1.9 kcal/mol), Y449 (1.0 kcal/mol), F456 (1.3 kcal/mol), F486 (2.1 kcal/mol), Y489 (1.8 kcal/mol), Q498 (1.2 kcal/mol), and Y505 (1.7 kcal/mol) (**Figure 3cd**). As was the case with the initial hot-spot prediction, the majority of the residues predicted to stabilize the complex are on the α1-helix (D30, K31, H34, D38, Y41). As in the KFC-2 hot-spot prediction, K353, which is found in the β3-β4 loop that flanks the α1-helix, was predicted as a stabilizing residue in the complex. Tyrosine 83, in the α2-helix (**Figure 1b, Figure 3c-d**), was also predicted as a contributor to complex stability. Predicted changes in binding free energies for all computational mutations, for both PMI

/ MDM2 and SARS-CoV-2 / ACE2, are available in the Supporting Information (**Figure S2**).

### Electrostatics and Topology at the Binding Interface

We used the Adaptive Poisson-Botzmann Solver (APBS)^16^ to generate electrostatic potential maps for the solvent-exposed surfaces of our positive control PPI (PMI/MDM2) and the SARS-CoV-2 RBD complex. As seen in **Figure 4a-b**, the positive control interaction is consistent with many PPIs, the driving factor for the PMI/MDM2 interaction is burying large, hydrophobic residues (most notably F3, W7, and L10) into a well-defined, largely hydrophobic, binding pocket. This results in relatively large changes in Gibbs’ free energy, as a result of beneficial changes in entropy (desolvation) and enthalpy (e.g. van der Waals interactions, π-π stacking). The binding face of the SARS-CoV-2 RBD / ACE2 complex (**Figure 4a**) was characterized as well. The binding surface of ACE2 has a pronounced patch with increased hydrophobic character, relative to the remainder of the surface, which aligns with the α1-helix (**Figure 4d**). Likewise, SARS-CoV-2 RBD presents a relatively shallow binding pocket with hydrophobic character, into which the α1-helix rests (**Figure 4d**). In contrast to the PMI / MDM2 complex, neither the ACE2 or SARS-CoV-2 surface contain deep, largely hydrophobic binding pockets. Collectively, this supports our hot-spot and computational alanine scanning mutagenesis studies. Compared to PMI / MDM2, more residues participate in stabilizing the complex; however, the magnitude of predicted stabilization is lower for each residue in the SARS-CoV-2 / ACE2 complex, compared to PMI / MDM2. We hypothesize that this is likely due to decreased burying of hydrophobic residues in the former, and increased hydrogen bonding and/or ion-paired interactions to stabilize the complex, relative to the later.

**Figure 4.**
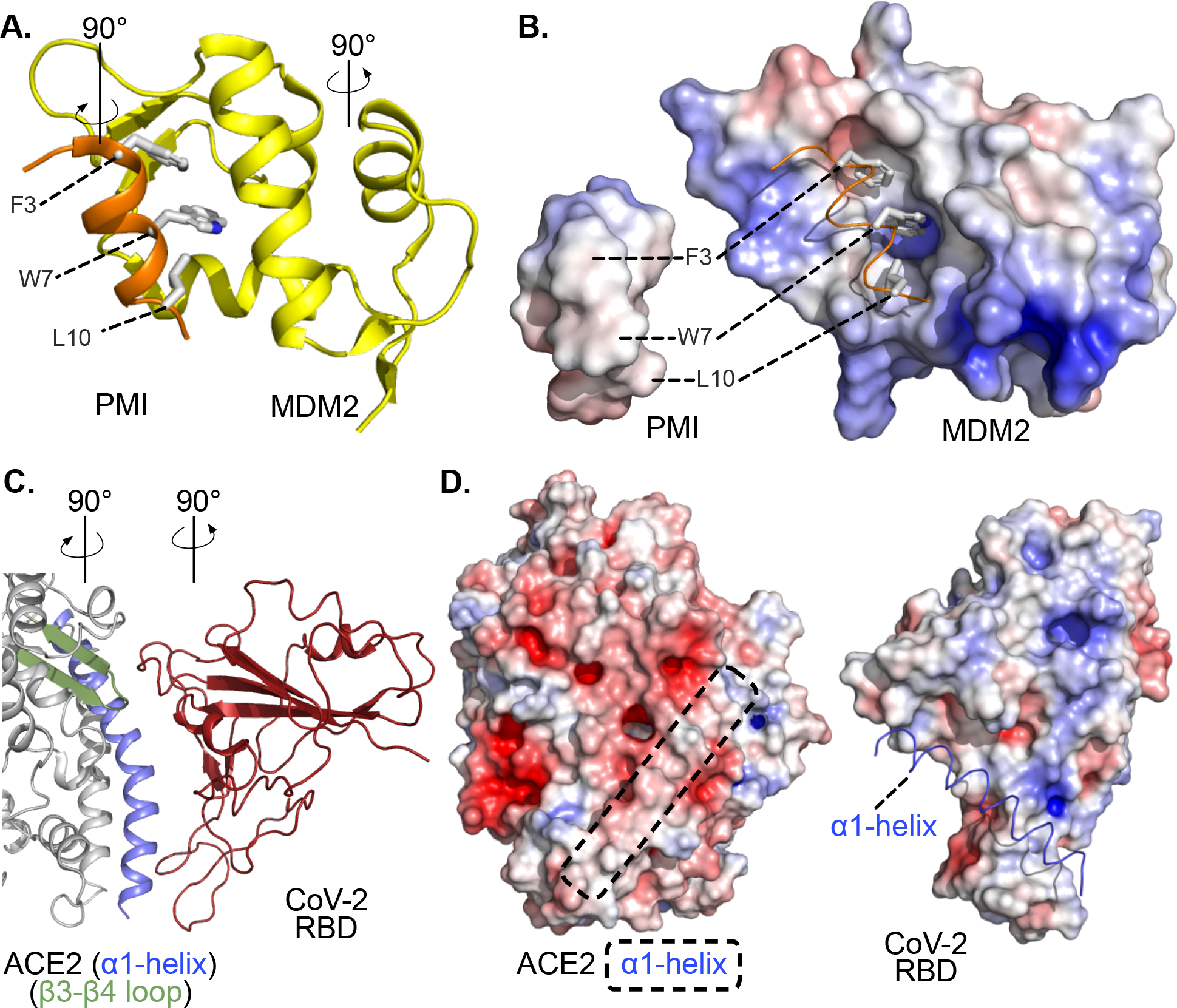
(**A**) Structure of the PMI / MDM2 complex. Hot spot residues (F3, W7, L10) are highlighed. (**B**) Electrostatic potential and topology of PMI and MDM2 surface areas that engage one another in the complex. (**C**) Strucuture of the ACE2 / SARS-CoV-2 RBD complex. α1-helix is highlighted in blue; β3-β4 loop is highlighted in green. (**D**) Electrostatic potential and topology of ACE2 and SARS-CoV-2 RBD surface areas that engage one another in the complex. Colored from −25 kT/e (red) to +25 kT/e (blue).

## Discussion

SARS-CoV-2, which causes Coronavirus-19 (COVID-19), is wreaking havoc on global economies and has been implicated in over 700,000 deaths worldwide. As an early step in the process of SARS-CoV-2 entry to the interior of mammalian cells (a necessary step in the virus lifecycle), SARS-CoV-2 RBD binds the extracellular protease domain (PD) of ACE2, a protein on the surface of cells in the lower respiratory tract. Inhibition of this Protein-Protein Interaction (PPI) is currently being evaluated as a therapeutic and/or prophylactic target. Recent structural studies have revealed key features of the SARS-CoV-2 RBD / ACE2 PD domain complex. Here, we conducted computational studies to predict hot-spot residues at the interface of this therapeutically-relevant PPI.

KFC-2 hot-spot prediction revealed five residues each on ACE2 and SARS-CoV-2 RBD that are predicted to stabilize the complex; computational alanine scanning with BAlaS predicted seven residues on each protein, which when mutated, significantly alter binding energy. There is good agreement between the two methods; both servers predicted ACE2 residues H34, D38, Y41, and K353, and SARS-CoV-2 RBD residues Y489, Q498, and Y505 as important for complex stability (**Figure 5**, residues highlighted with an asterisk).

**Figure 5.**
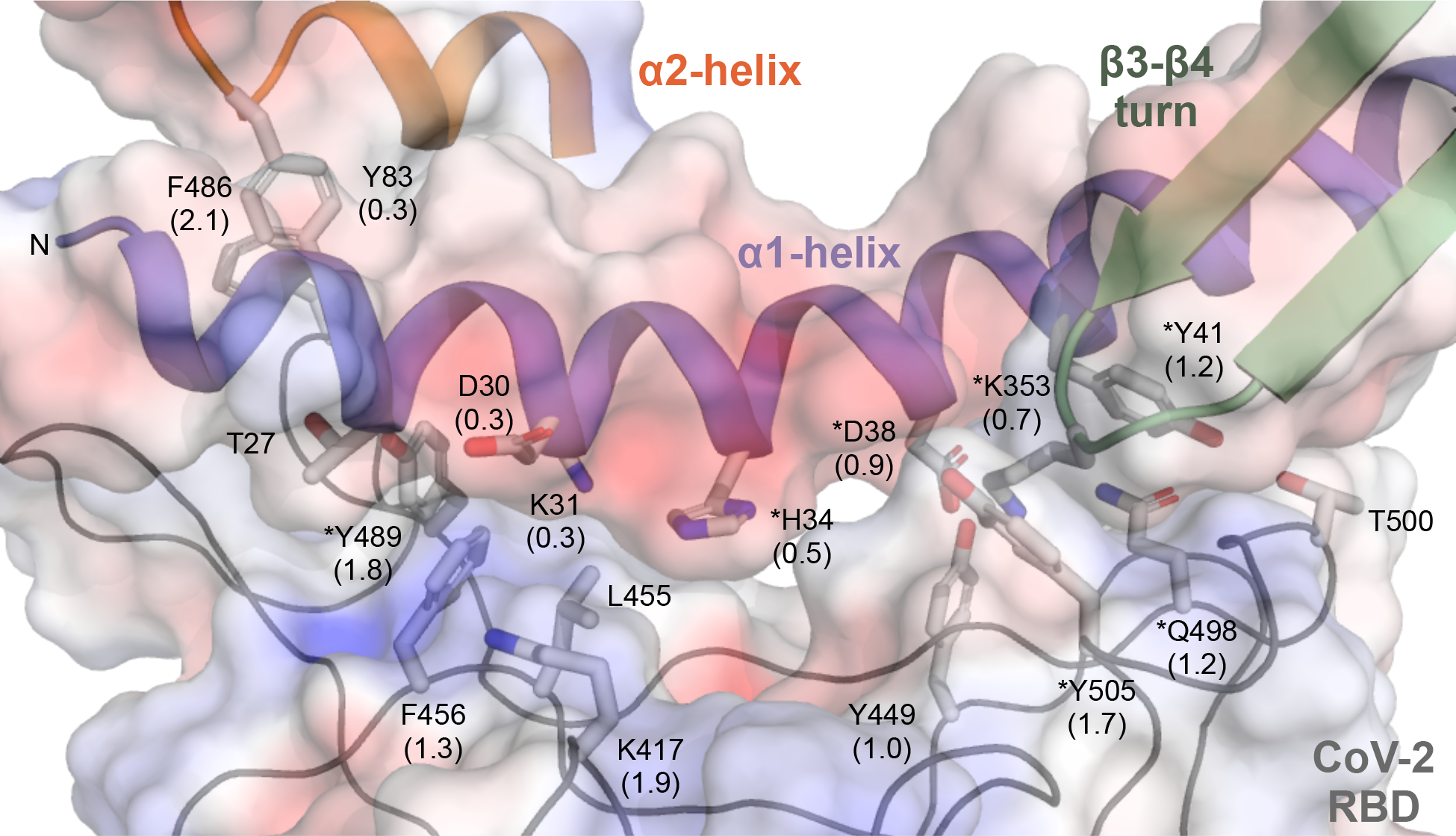
Residues on the α1-helix (blue), α-2 helix (orange), and β3-β4 loop (green) on ACE2 (top) and SARS-CoV-2 RBD (bottom) that were identified as stabilizing formation of a complex between these two proteins. Structures and selected residues are overlaid with the electrostatic potential map of protein surfaces at the binding interface. Values in parenthesis are predicted ΔΔG (kcal/mol) values obtained from computational alanine scanning mutagenesis. An asterisk (*) indicates that the residue was predicted by they KFC-2 hot-spot server and BAlaS computational alanine scanning server.

Interestingly, while the structure of SARS-CoV RBD aligns well with SARS-CoV-2 RBD, with a root mean squared deviation (RMSD) of 0.68 Å over 139 pairs of Cα atoms^8^, mutations exist, including interface residues predicted here to stabilize SARS-CoV-2 RBD interactions with ACE2. For example, Y484®Q498, L443®F456, N479®Q493, and L472®F486 mutations at equivalent positions of SARS-CoV RBD and SARS-CoV-2 RBD, respectively, are observed.^7, 8^ These mutations may change the affinity between SARS-CoV-2 for ACE2, relative to SARS-CoV, and likely alters the hot-spot geographies on the binding surface of SARS-CoV-2 RBD, compared to SARS-CoV.

## Acknowledgements

We acknowledge Delaware State University for institutional support and the Delaware INBRE program, which is supported by a grant from the National Institute of General Medical Sciences (NIGMS, P20 GM103446) from the National Institutes of Health and the State of Delaware.

