## Supporting Information for "Computational Hot-Spot Analysis of the SARS-CoV-2 Receptor Binding Domain / ACE2 Complex"

**Authors:** Pedro A. Rosario[1] and Brian R. McNaughton[2,*]
**Affiliations:**
[1]PEMaCS Division, Physics & Engineering Program, Delaware State University, Dover, DE 19901, USA
[2]Delaware Institute for Science & Technology, Delaware State University, Dover, DE 19901, USA

### Supporting Information and Figures

##### *1. Hot-spot prediction*

Hot spot prediction was performed using KFC-2 () as previously described. PDB codes (3eqs for PMI/MDM2 or 6m17 for SARS-CoV-2 RBD/ACE2) were loaded to the server. Chains A (MDM2) and B (PMI) were used in 3eqs; chains D (ACE2 PD) and F (SARS-CoV-2 RBD) were used in 6m17. Data is shown in **Figure S1**.

##### *2. Computational alanine scanning mutagenesis*

Computational mutagenesis was performed using the BAlaS server (), as previously described. PDB files (3eqs for PMI/MDM2 or 6m17 for SARS-CoV-2 RBD/ACE2) were loaded to the server. Chains A and B were used in 3eqs; chains D and F were used in 6m17. Distance cut-off was set at 13 Å; change in free energy cut-off was set at 4 kJ/mol. Energies were converted to kcal/mol for representation in this manuscript. Data is shown in **Figure S2**.

##### *3. Electrostatics*

Electrostatic potential maps were prepared in PyMol using the APBS electrostatics plugin.

**A**

KFC2 Hot Spot Prediction Server @mitchell-lab.org from Tue, 28 Jul 2020 19:43:40 EDT
JobId: 20e2c6e32a8b7d46c01063835f6722fa   JobName: 3   ChainSet1: A   ChainSet2: B

| Chain | Res | Num | KFC2-A Class | KFC2-A Conf | KFC2-B Class | KFC2-B Conf | ConSurf Class | ConSu Value | Rosetta Class | Roset DDG | Exper Class | Exper Value |
|---|---|---|---|---|---|---|---|---|---|---|---|---|
| A | LYS | 51 | ------- | -2.42 | ------- | -0.97 | ------- | --- | ------- | --- | ------- | --- |
| A | LEU | 54 | Hotspot | 0.64 | Hotspot | 0.20 | ------- | --- | ------- | --- | ------- | --- |
| A | PHE | 55 | ------- | -1.99 | ------- | -0.92 | ------- | --- | ------- | --- | ------- | --- |
| A | LEU | 57 | ------- | -0.18 | Hotspot | 0.07 | ------- | --- | ------- | --- | ------- | --- |
| A | GLY | 58 | ------- | -0.84 | ------- | -0.66 | ------- | --- | ------- | --- | ------- | --- |
| A | ILE | 61 | ------- | -0.07 | Hotspot | 0.06 | ------- | --- | ------- | --- | ------- | --- |
| A | MET | 62 | ------- | -1.10 | ------- | -0.83 | ------- | --- | ------- | --- | ------- | --- |
| A | TYR | 67 | ------- | -1.76 | ------- | -0.80 | ------- | --- | ------- | --- | ------- | --- |
| A | GLN | 71 | ------- | -2.83 | ------- | -0.94 | ------- | --- | ------- | --- | ------- | --- |
| A | GLN | 72 | ------- | -0.30 | ------- | -0.43 | ------- | --- | ------- | --- | ------- | --- |
| A | HIS | 73 | ------- | -0.64 | ------- | -0.63 | ------- | --- | ------- | --- | ------- | --- |
| A | VAL | 75 | Hotspot | 0.56 | ------- | -0.05 | ------- | --- | ------- | --- | ------- | --- |
| A | PHE | 86 | ------- | -2.81 | ------- | -0.88 | ------- | --- | ------- | --- | ------- | --- |
| A | PHE | 91 | ------- | -1.04 | ------- | -0.78 | ------- | --- | ------- | --- | ------- | --- |
| A | VAL | 93 | Hotspot | 1.33 | Hotspot | 0.06 | ------- | --- | ------- | --- | ------- | --- |
| A | LYS | 94 | ------- | -2.31 | ------- | -0.92 | ------- | --- | ------- | --- | ------- | --- |
| A | HIS | 96 | ------- | -0.95 | ------- | -0.78 | ------- | --- | ------- | --- | ------- | --- |
| A | ILE | 99 | Hotspot | 0.78 | Hotspot | 0.19 | ------- | --- | ------- | --- | ------- | --- |
| A | TYR | 100 | ------- | -1.40 | ------- | -0.78 | ------- | --- | ------- | --- | ------- | --- |
| A | MET | 102 | ------- | -3.16 | ------- | -0.86 | ------- | --- | ------- | --- | ------- | --- |
| B | THR | 1 | ------- | -2.25 | ------- | -0.94 | ------- | --- | ------- | --- | ------- | --- |
| B | SER | 2 | ------- | -1.72 | ------- | -0.90 | ------- | --- | ------- | --- | ------- | --- |
| B | PHE | 3 | Hotspot | 1.55 | Hotspot | 0.23 | ------- | --- | ------- | --- | ------- | --- |
| B | ALA | 4 | ------- | -1.01 | ------- | -0.74 | ------- | --- | ------- | --- | ------- | --- |
| B | TYR | 6 | Hotspot | 0.59 | ------- | -0.00 | ------- | --- | ------- | --- | ------- | --- |
| B | TRP | 7 | Hotspot | 1.04 | Hotspot | 0.28 | ------- | --- | ------- | --- | ------- | --- |
| B | ASN | 8 | ------- | -1.59 | ------- | -0.68 | ------- | --- | ------- | --- | ------- | --- |
| B | LEU | 9 | ------- | -1.77 | ------- | -0.78 | ------- | --- | ------- | --- | ------- | --- |
| B | LEU | 10 | Hotspot | 1.01 | Hotspot | 0.08 | ------- | --- | ------- | --- | ------- | --- |
| B | SER | 11 | ------- | -0.72 | ------- | -0.86 | ------- | --- | ------- | --- | ------- | --- |

**B**

JobId: 5613f8aa3bbf07c1a5f5abb9259f2069   JobName: 2   ChainSet1: D   ChainSet2: F

| Chain | Res | Num | KFC2-A Class | KFC2-A Conf | KFC2-B Class | KFC2-B Conf | ConSurf Class | ConSu Value | Rosetta Class | Roset DDG | Exper Class | Exper Value |
|---|---|---|---|---|---|---|---|---|---|---|---|---|
| D | ILE | 21 | ------- | -3.12 | ------- | -0.93 | ------- | --- | ------- | --- | ------- | --- |
| D | GLN | 24 | ------- | -0.13 | ------- | -0.36 | ------- | --- | ------- | --- | ------- | --- |
| D | THR | 27 | Hotspot | 0.31 | ------- | -0.67 | ------- | --- | ------- | --- | ------- | --- |
| D | PHE | 28 | ------- | -1.32 | ------- | -0.57 | ------- | --- | ------- | --- | ------- | --- |
| D | ASP | 30 | ------- | -1.85 | ------- | -1.00 | ------- | --- | ------- | --- | ------- | --- |
| D | LYS | 31 | ------- | -0.49 | ------- | -0.44 | ------- | --- | ------- | --- | ------- | --- |
| D | HIS | 34 | Hotspot | 0.27 | ------- | -0.57 | ------- | --- | ------- | --- | ------- | --- |
| D | GLU | 35 | ------- | -1.68 | ------- | -0.94 | ------- | --- | ------- | --- | ------- | --- |
| D | GLU | 37 | ------- | -1.42 | ------- | -0.83 | ------- | --- | ------- | --- | ------- | --- |
| D | ASP | 38 | Hotspot | 0.45 | ------- | -0.54 | ------- | --- | ------- | --- | ------- | --- |
| D | TYR | 41 | Hotspot | 0.93 | Hotspot | 0.35 | ------- | --- | ------- | --- | ------- | --- |
| D | GLN | 42 | ------- | -2.19 | ------- | -0.99 | ------- | --- | ------- | --- | ------- | --- |
| D | LEU | 45 | ------- | -1.46 | ------- | -0.90 | ------- | --- | ------- | --- | ------- | --- |
| D | LEU | 79 | ------- | -2.11 | ------- | -0.92 | ------- | --- | ------- | --- | ------- | --- |
| D | MET | 82 | ------- | -2.84 | ------- | -0.91 | ------- | --- | ------- | --- | ------- | --- |
| D | TYR | 83 | ------- | -0.82 | ------- | -0.29 | ------- | --- | ------- | --- | ------- | --- |
| D | THR | 324 | ------- | -1.90 | ------- | -0.95 | ------- | --- | ------- | --- | ------- | --- |
| D | ASN | 330 | ------- | -1.87 | ------- | -0.96 | ------- | --- | ------- | --- | ------- | --- |
| D | LYS | 353 | Hotspot | 1.21 | Hotspot | 0.10 | ------- | --- | ------- | --- | ------- | --- |
| D | GLY | 354 | ------- | -1.32 | ------- | -0.74 | ------- | --- | ------- | --- | ------- | --- |
| D | ASP | 355 | ------- | -0.09 | ------- | -0.38 | ------- | --- | ------- | --- | ------- | --- |
| D | ARG | 357 | ------- | -1.16 | ------- | -0.31 | ------- | --- | ------- | --- | ------- | --- |
| D | ALA | 386 | ------- | -2.35 | ------- | -0.82 | ------- | --- | ------- | --- | ------- | --- |
| D | ARG | 393 | ------- | -2.98 | ------- | -0.77 | ------- | --- | ------- | --- | ------- | --- |
| F | ARG | 403 | ------- | -1.86 | ------- | -0.73 | ------- | --- | ------- | --- | ------- | --- |
| F | LYS | 417 | ------- | -2.84 | ------- | -0.93 | ------- | --- | ------- | --- | ------- | --- |
| F | VAL | 445 | ------- | -2.10 | ------- | -0.93 | ------- | --- | ------- | --- | ------- | --- |
| F | GLY | 446 | ------- | -2.30 | ------- | -0.74 | ------- | --- | ------- | --- | ------- | --- |
| F | TYR | 449 | ------- | -1.82 | ------- | -0.83 | ------- | --- | ------- | --- | ------- | --- |
| F | TYR | 453 | ------- | -1.27 | ------- | -0.21 | ------- | --- | ------- | --- | ------- | --- |
| F | LEU | 455 | Hotspot | 0.29 | Hotspot | 0.08 | ------- | --- | ------- | --- | ------- | --- |
| F | PHE | 456 | ------- | -0.27 | ------- | -0.04 | ------- | --- | ------- | --- | ------- | --- |
| F | TYR | 473 | ------- | -1.74 | ------- | -0.82 | ------- | --- | ------- | --- | ------- | --- |
| F | ALA | 475 | ------- | -1.62 | ------- | -0.84 | ------- | --- | ------- | --- | ------- | --- |
| F | GLY | 476 | ------- | -1.81 | ------- | -0.76 | ------- | --- | ------- | --- | ------- | --- |
| F | GLY | 485 | ------- | -1.97 | ------- | -0.75 | ------- | --- | ------- | --- | ------- | --- |
| F | PHE | 486 | ------- | -0.68 | ------- | -0.64 | ------- | --- | ------- | --- | ------- | --- |
| F | ASN | 487 | ------- | -1.81 | ------- | -0.72 | ------- | --- | ------- | --- | ------- | --- |
| F | TYR | 489 | Hotspot | 0.73 | Hotspot | 0.04 | ------- | --- | ------- | --- | ------- | --- |
| F | PHE | 490 | ------- | -2.91 | ------- | -0.87 | ------- | --- | ------- | --- | ------- | --- |
| F | GLN | 493 | ------- | -0.32 | ------- | -0.39 | ------- | --- | ------- | --- | ------- | --- |
| F | TYR | 495 | ------- | -1.43 | ------- | -0.70 | ------- | --- | ------- | --- | ------- | --- |
| F | GLY | 496 | ------- | -0.85 | ------- | -0.68 | ------- | --- | ------- | --- | ------- | --- |
| F | GLN | 498 | Hotspot | 0.76 | Hotspot | 0.09 | ------- | --- | ------- | --- | ------- | --- |
| F | THR | 500 | Hotspot | 0.41 | ------- | -0.65 | ------- | --- | ------- | --- | ------- | --- |
| F | ASN | 501 | ------- | -0.28 | ------- | -0.39 | ------- | --- | ------- | --- | ------- | --- |
| F | GLY | 502 | ------- | -0.53 | ------- | -0.66 | ------- | --- | ------- | --- | ------- | --- |
| F | VAL | 503 | ------- | -2.08 | ------- | -1.01 | ------- | --- | ------- | --- | ------- | --- |
| F | TYR | 505 | Hotspot | 0.08 | ------- | -0.20 | ------- | --- | ------- | --- | ------- | --- |

**Figure S1.** KFC-2 data for (**A**) PMI / MDM2 (PDB: 3eqs) and (**B**) SARS-CoV-2 RBD / ACE2 (PDB: 6m17)

**A**

| Chain | Residue | Amino Acid | ΔΔG (kJ/mol) |
|---|---|---|---|
| A | 54 | LEU | 4.6496 |
| A | 67 | TYR | 5.2389 |
| A | 72 | GLN | 7.1022 |
| A | 93 | VAL | 6.2081 |
| A | 96 | HIS | 4.4006 |

| Chain | Residue | Amino Acid | ΔΔG (kJ/mol) |
|---|---|---|---|
| B | 3 | PHE | 19.3057 |
| B | 6 | TYR | 10.1194 |
| B | 7 | TRP | 21.9557 |
| B | 10 | LEU | 8.7882 |

**B**

| Chain | Residue | Amino Acid | ΔΔG (kJ/mol) |
|---|---|---|---|
| D | 30 | ASP | 7.9964 |
| D | 31 | LYS | 3.723 |
| D | 34 | HIS | 3.9799 |
| D | 38 | ASP | 3.1237 |
| D | 41 | TYR | 8.813 |
| D | 83 | TYR | 4.4579 |
| D | 353 | LYS | 5.8613 |

| Chain | Residue | Amino Acid | ΔΔG (kJ/mol) |
|---|---|---|---|
| F | 417 | LYS | 7.9416 |
| F | 449 | TYR | 4.1881 |
| F | 456 | PHE | 5.311 |
| F | 486 | PHE | 8.7995 |
| F | 489 | TYR | 7.7264 |
| F | 498 | GLN | 4.8124 |
| F | 505 | TYR | 7.0366 |

**Figure S2.** Data obtained from the BAlaS server for (**A**) PMI/MDM2 and (**B**) ACE2/SARS-CoV-2 RBD
